# Modeling the underlying environmental factors of milky sea case and luminous bacteria presence in Java Southern Sea in 2019

**DOI:** 10.1101/2021.09.10.459852

**Authors:** Andri Wibowo

## Abstract

The milky sea is one of the unique natural phenomena caused by the presence of luminous *Vibrio* bacteria in marine ecosystems. Recently a milky sea has been reported frequently included in the Java Southern Sea. Simultaneously, numerous remote sensing based approaches have been developed to detect the presence of luminous bacteria and the milky sea. Despite this state of the art, the information of detrimental factors of the marine bioluminescence was still limited. Then this research aims to model the underlying environmental factors causing the milky sea and luminous bacteria presence in the Java Southern Sea in 2019. The remote sensing assessment for the period of July 29-August 6, 2019 shows that the magnitude of bioluminescence measured in radiance was having a maximum value of 255 nanoW/cm^2^/sr and an average of 107 nanoW/cm^2^/sr/day (95%CI: 71.9 to 142 nanoW/cm^2^/sr/day). The milky sea size increased and reached its peak with a size of 44,124 km^2^ and then declined. The average milky sea size was 37,942 km^2^ (95% CI: 33,400 to 42,500 km^2^) and increased with average rate of 16.01% (95%CI: 5.41% to 26.66%). While Akaike Information Criterion (AIC) indicates that the best model to infer the relationship of bacterial bioluminescence with its environmental factors contained Chlorophyll a followed by sea surface temperature factors with AIC_c_ values of 101.16 (AIC_weight_: 0.50) and 101.95 (AIC_weight_: 0.34). This indicates that low temperature and high plankton cells is the limiting factors of the bacterial bioluminescence.

## Introduction

In nature, organism mainly microorganism has the ability to emit the light. This ability is known as bioluminescence. This bioluminescence was also observed in aquatic ecosystems including in marine ecosystems. Bioluminescence is a result of the biochemistry reaction of organisms involving the oxidation of an aliphatic aldehyde by a reduced flavin mononucleotide. The products of this oxidation reaction include an oxidized flavin mononucleotide, a fatty acid chain, and energy in the form of a blue-green visible light. The bioluminescence process was as follows: FMNH_2_ + O_2_ + RCHO → FMN + RCOOH + H_2_O + light (Fisher et al. 1996, Willes et al. 2005, Gregor et al. 2018). The purpose of bioluminescence is to attract prey or even more complicated. In bacteria, bioluminescence functions to attract and lure other organisms to prey on bacteria (Zarubin et al. 2012).

There are wide ranges of marine microorganisms that can emit lights, including bacteria. In bacteria, bioluminescence occurs as a continuous glow in the presence of oxygen at cell concentrations exceeding quorum-sensing levels. Luminous bacteria were occurring as free-living microorganisms in seawater, in symbiotic associations with marine organisms (most notably fish and squids), as saprophytes on suspended organic material such as marine snow, as a major component of fecal pellets, and as parasites on crustaceans (Nelson & Hastings 1979, Ruby et al. 1979, Dworkin et al. 2006, Andrew et al. 1984). Among several bioluminescent bacteria species inhabiting marine ecosystems, there are species of interest including members of *Vibrio* genera. The bacterial bioluminescence is characterized by its continuous light emission that can persist for many days under appropriate conditions.

A bioluminescence activity if happens in the sea will be appeared as milky sea due to the layer of whitish light expanding more than 1 km^2^ in the surface of the water. Recently the detection of the milky sea and bacterial bioluminescence has received attentions. The study was focused on the development of low light sensor that can detect bioluminescence. The most prominent studies on luminous detection were using satellite observation and Visible Infrared Imager Radiometer Suite. Using this method, Miller et al. (2005) has identified bacterial bioluminescence in the northwestern Indian Ocean spanning over an area sizing 15,400 km^2^ and glow over 3 consecutive nights with emission spectra give peak values at 490 nm and half-bandwidths of 70 nm. *Vibrio fischeri* strains whose emission spectra are representative of the luminous bacteria species thought to be responsible for milky seas. Recently, bioluminescence has been identified in several locations including in the Java Southern Sea in recent 2019 (Miller et al. 2021). Despite current detection of milky sea in the Java Southern Sea, information on determinant environmental factors causing the presence of milky sea and bacterial bioluminescence in Java Southern Sea is still limited. Then this study aims to model the underlying environmental factors of the milky sea and luminous bacteria presences in the Java Southern Sea in 2019.

## Methodology

### Study area

The observed area was the Java Southern Sea located on the South of Java Island. The observation period was from 29 July 2019 to 6 August 2019. During this period, several parameters were retrieved using satellite imagery including the presence of the milky sea measured in, sea surface temperature/SST (°C), and Chlorophyll a contents (mg/m^3^).

### Bioluminescence detection

The presence of milky sea and bacterial luminescence were measured as radiance denoted as nanoW/cm^2^/sr. The radiance was measured followed Baugh et al. (2013). The platforms to detect bioluminescence were the Day/Night Band (DNB) parts of the Visible Infrared Imaging Radiometer Suite carried on National Oceanic and Atmospheric Administration operational satellites. This satellite covers regions between latitude of −65° South and +75° North and produce versatile nighttime images. Monthly composites of those 16-bit images processed by the National Centers for Environmental Information, National Oceanic and Atmospheric Administration were used to detect luminescence with spatial resolution of 15 arc second, equivalent to approximately 0.5 km on the equator (Hillger et al. 2014, Román et al. 2018, Priyatikanto et al. 2019).

### Sea surface temperature

The sea surface temperature (SST) model of Java Southern Sea for a period of 29 July-6 August 2019 was developed using a method followed Martin et al. (2012). The SST is developed upon the Group for High Resolution Sea Surface Temperature SST used wavelets as basic functions in an optimal interpolation approach on a global 0.011 degree grid. This method is based upon observations from several instruments including the NASA Advanced Microwave Scanning Radiometer-EOS and the Moderate Resolution Imaging Spectroradiometer on the NASA Aqua and Terra platforms.

### Chlorophyll a content

Chlorophyll a contents in Java Southern Sea was measured and mapped based on Terra Moderate Resolution Imaging Spectroradiometer and method followed Ghanea et al. (2015). Terra Moderate Resolution Imaging Spectroradiometer with a spatial resolution of 1 km were downloaded from NASA data archive and processed to L3 products. The data were composites from a period of 29 July-6 August 2019 that has been processed according to the following algorithm:

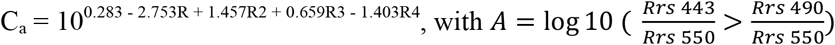

Where C_a_ is the concentration of Chlorophyll a denoted as mg/m^3^, R is the reflectance ratio, and Rrs is the remote sensing reflectance (Alianto & Hamuna 2020).

### Statistical analysis

Statistical analysis aims to assess the underlying factors causing the presence of bioluminescence. The method used was Principal Component Analysis to determine the correlations and best model of bioluminescence with the underlying factors including SST and Chlorophyll a. The models were validated using Akaike Information Criterion (AIC).

## Results and Discussions

### Bioluminescence detection

The results of bacterial bioluminescence detection using Visible Infrared Imaging Radiometer Suite sensors for a period of July 29-August 6 were available in Figure 1. It is obvious that the appearance of milk sea and bioluminescence in the Java Southern Sea was started on July 29 appeared in West then it was gradually increased in radiance and size (Figure 2). The magnitude of bioluminescence measured in radiance was having a maximum value of 255 nanoW/cm^2^/sr and an average of 107 nanoW/cm^2^/sr/day (95%CI: 71.9 to 142 nanoW/cm^2^/sr/day). The bioluminescence was declining after 9 days or on August 6. The size of milky sea increased and reached its peak with a size of 44,124 km^2^ and then declined. The average milky sea size was 37,942 km^2^ (95% CI: 33,400 to 42,500 km^2^). For 9 days from July 29 to August 6, the size of the milky sea in the Java Southern Sea increased on an average of 16.01% (95%CI: 5.41% to 26.66%). On August 6, East parts of milky sea were reduced.

**Figure 1.**
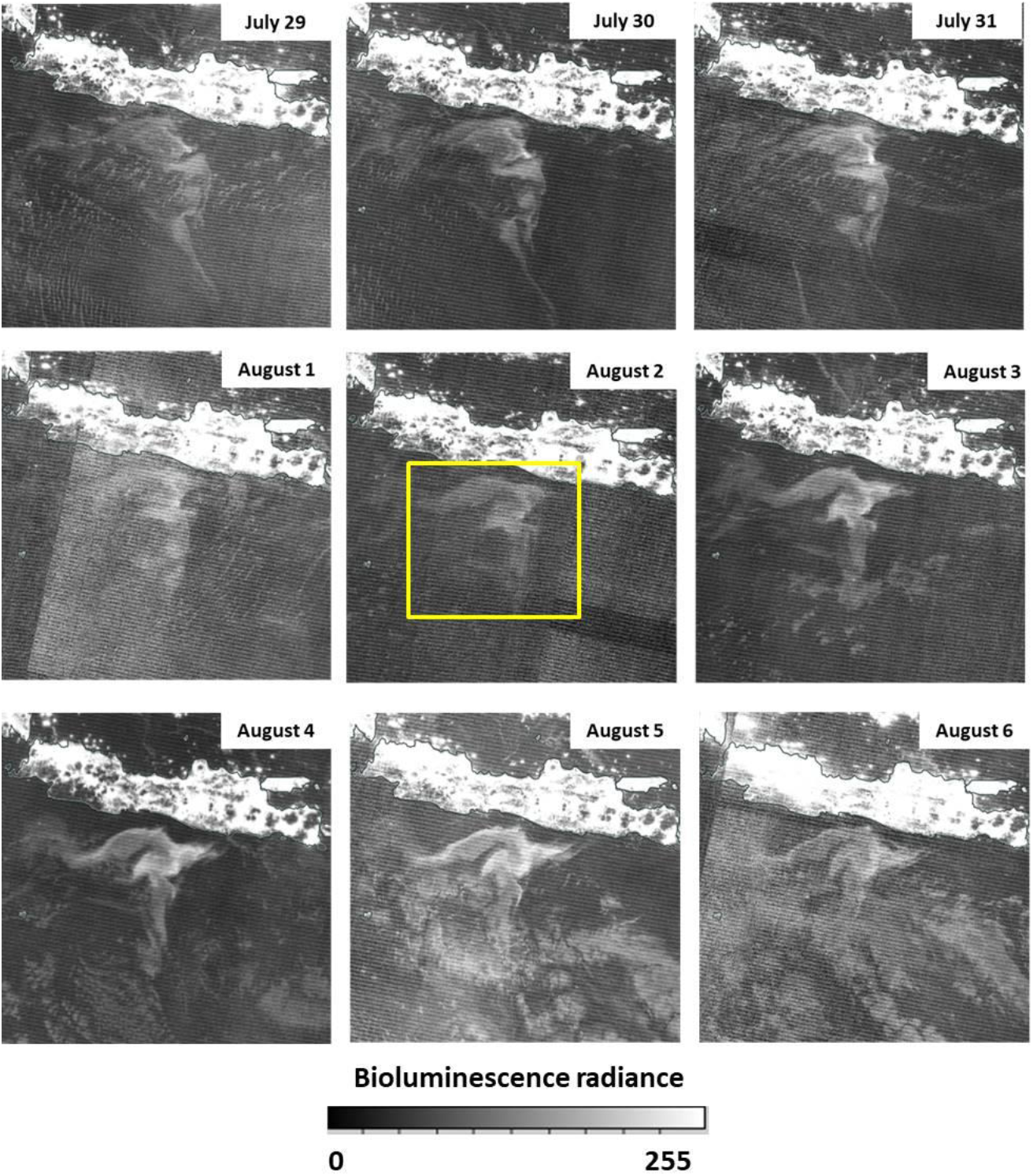
Patterns and appearances of milky sea and bacterial bioluminescence radiance (yellow square and denoted as nanoW/cm^2^/sr) in the Java Southern Sea for period of July 29-August 6, 2019 (Source: Visible Infrared Imaging Radiometer Suite).

**Figure 2.**
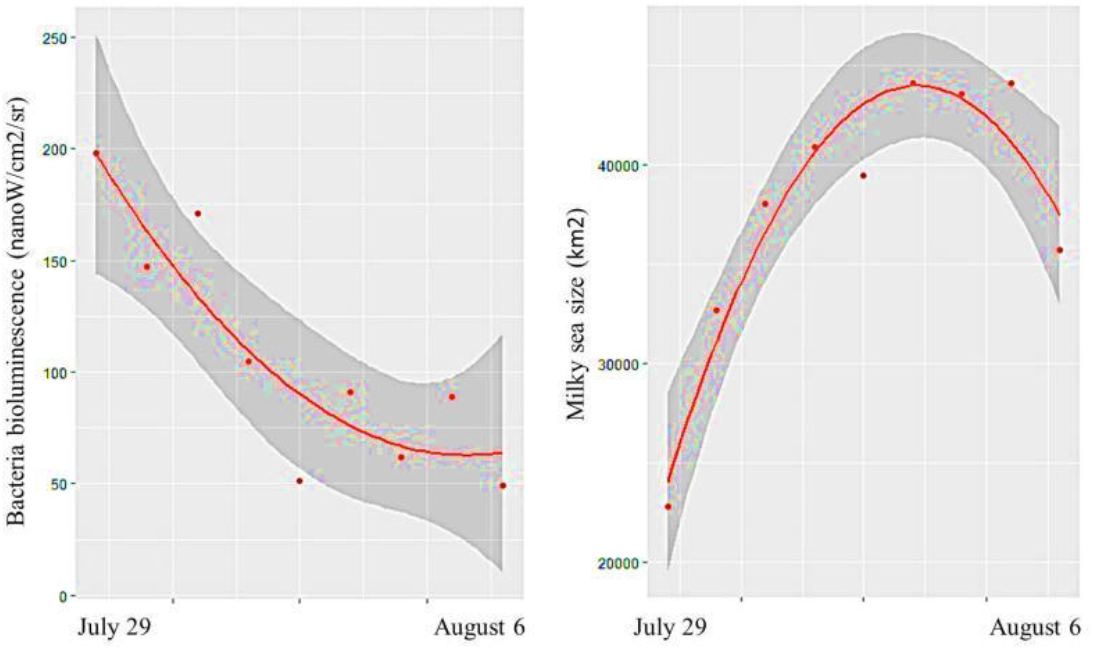
Trends with 95%CI of bacterial bioluminescence radiance (left, nanoW/cm^2^/sr) and size of milky sea (right, km^2^) in the Java Southern Sea for period of July 29-August 6, 2019.

### Sea surface temperature

The sea surface temperature (SST) of Java Southern Sea has an average of 24.6 °C (95%CI: 24.2 to 25 °C). It is apparent that the water of the Java Southern Sea was warmer whereas there were parts of the Java Southern Sea that has lower SST. This area was observed in the central of Java Southern Sea (Figure 3). In comparison to milky sea presences, the bioluminescence radiance was not presented in the water that has lower SST. Whereas, bioluminescence radiance was observed high in parts of the Java Southern Sea that have high SST. This fact is also supported by the linear trends of bioluminescence radiance with SST (Figure 4). It was apparent that on August 6 the size of water that has lower temperature increased and expanded to the West and reduced the size of the milky sea. The average Chlorophyll a in Java Southern Sea was 0.626 mg/m^3^ (95%CI: 0.528 to 0.712 mg/m^3^) (Figure 5). Bioluminescence radiance was observed high in parts of the Java Southern Sea that has low Chlorophyll a. This means that increase in Chlorophyll a will reduce the bioluminescence radiance.

**Figure 3.**
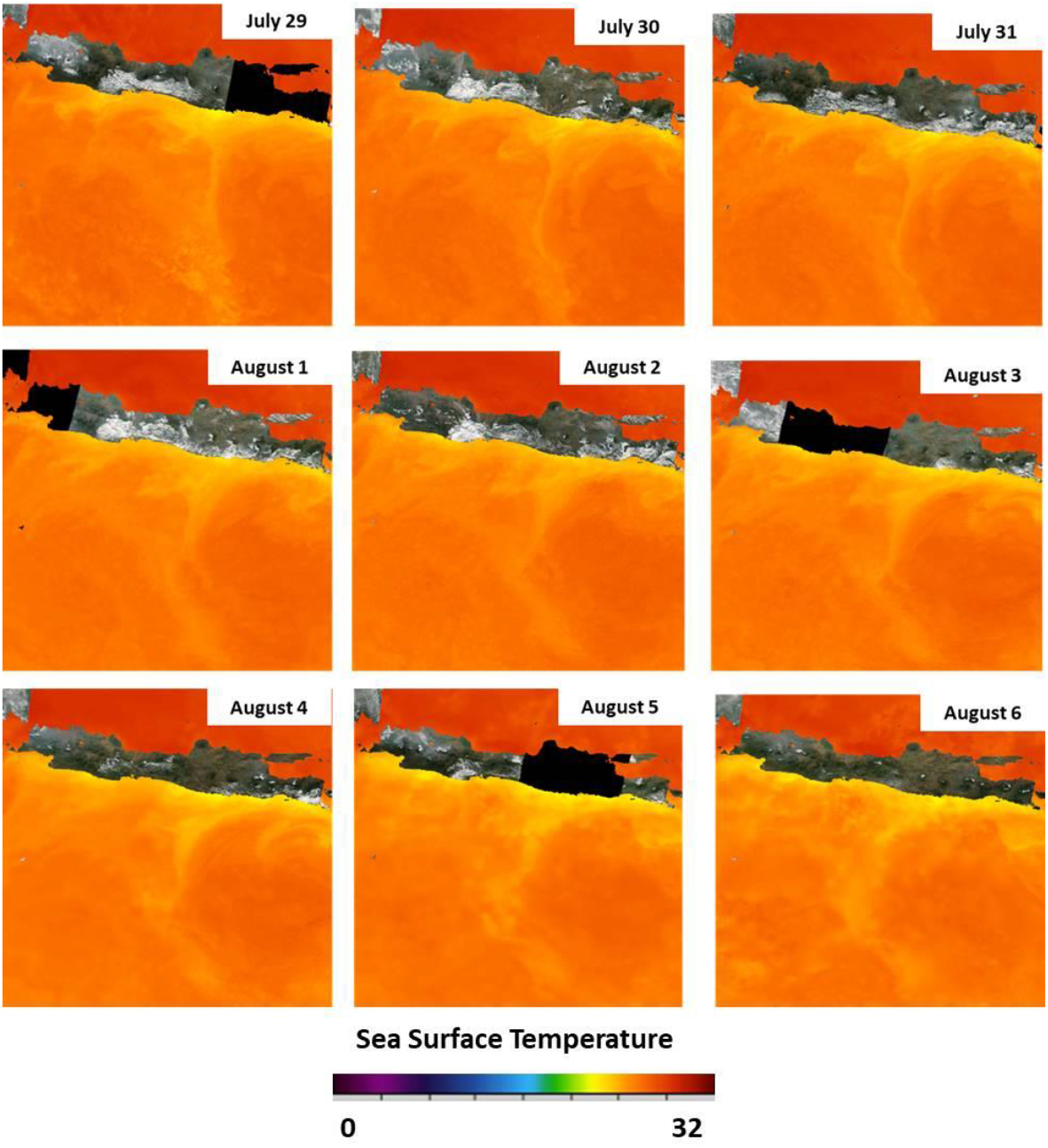
Patterns of sea surface temperature/SST (^0^C) in the Java Southern Sea for period of July 29-August 6, 2019 (Source: the Group for High Resolution Sea Surface Temperature).

**Figure 4.**
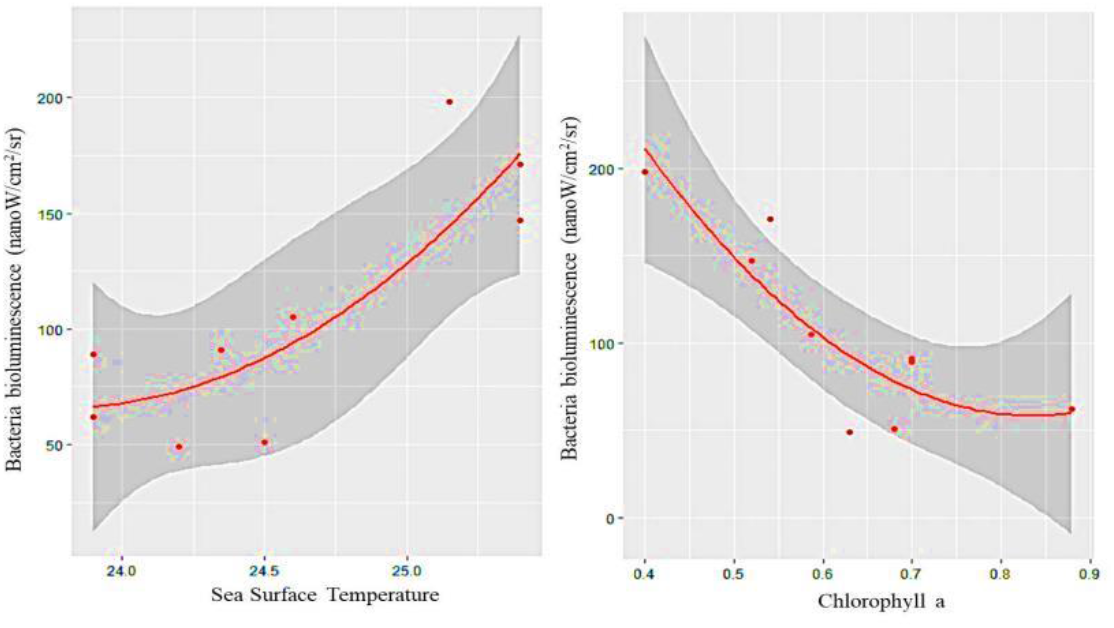
Trends with 95%CI of bacterial bioluminescence radiance (nanoW/cm^2^/sr) with SST (left, °C) and Chlorophyll a (right, mg/m^3^) in the Java Southern Sea for period of July 29-August 6, 2019.

**Figure 5.**
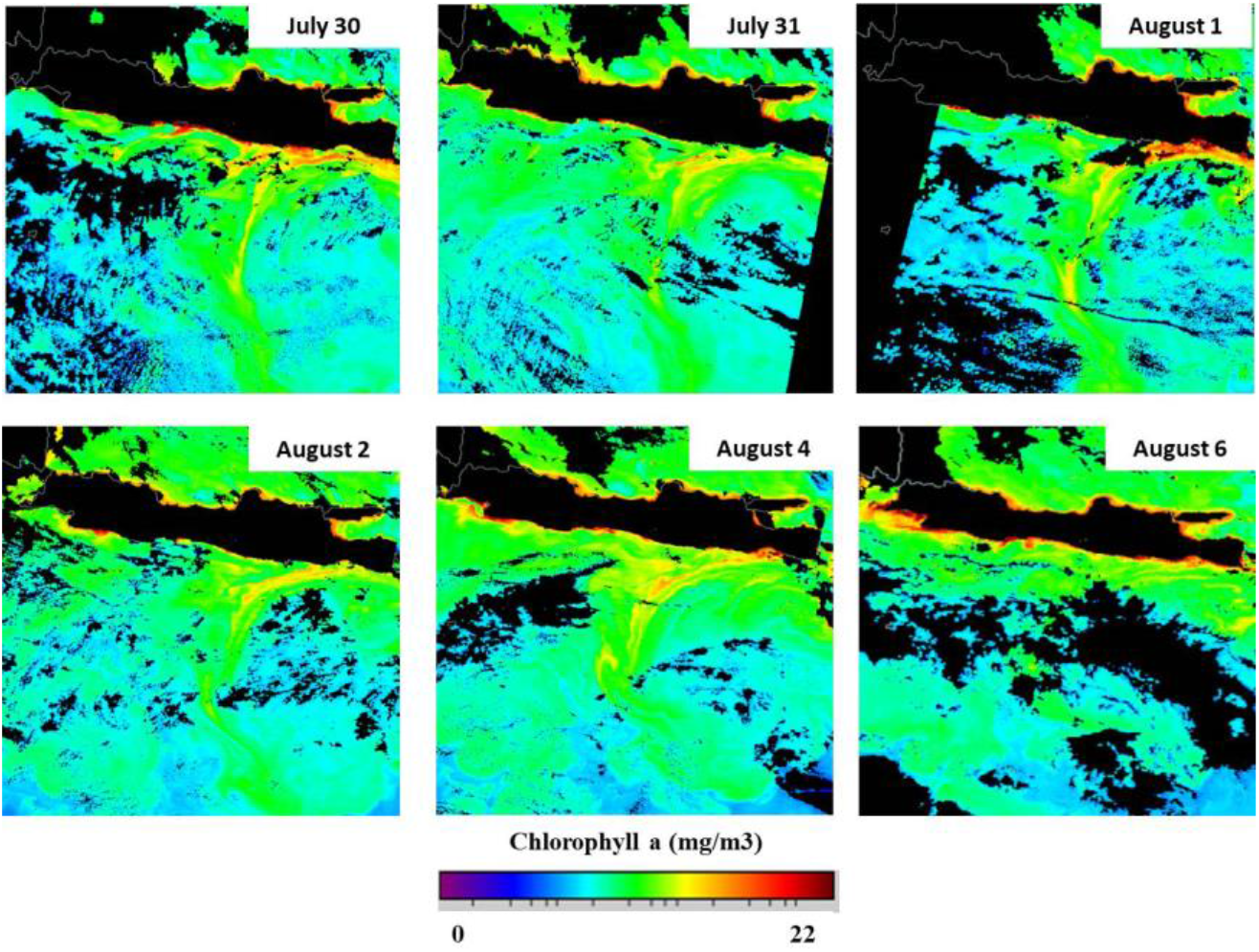
Patterns of Chlorophyll a (mg/m^3^) in the Java Southern Sea for period of July 30 - August 6, 2019 (Source: the Group for High Resolution Sea Surface Temperature).

**Figure 6.**
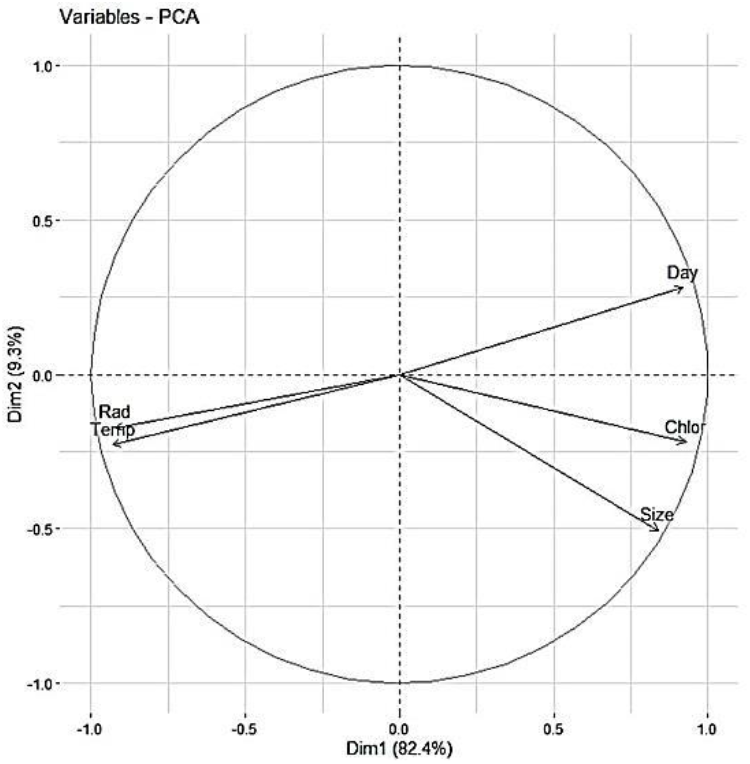
PCA of bacterial bioluminescence (Rad), milky sea size (Size), with Chlorophyll a (Chlor), temperature (Temp), and days (Day) in the Java Southern Sea for period of July 30 - August 6, 2019.

**Table 1.** Akaike Information Criterion (AIC) attributes for selected models of bacterial bioluminescence (B), Chlorophyll a (C), temperature (T), and days (D).

| Model | K | AIC<sub>c</sub> | ΔAIC | AIC<sub>weight</sub> | Cum.-weight | Log Likelihood |
| --- | --- | --- | --- | --- | --- | --- |
| B ~ C + D | 4 | 101.16* | 0.00 | 0.50 | 0.50 | -41.58 |
| B ~ T + D | 4 | 101.95* | 0.79 | 0.34 | 0.84 | -41.98 |
| B ~ C + T | 4 | 103.42 | 2.26 | 0.16 | 1.00 | -42.71 |
| B ~ C + D + T | 5 | 113.12 | 11.95 | 0.00 | 1.00 | -41.56 |

\*Best model

### Model of milky sea and bacterial bioluminescence

PCA has confirmed that the high temperature and low Chlorophyll a were correlated with high bacterial bioluminescence. While Akaike Information Criterion (AIC) indicates that the best model to infer the relationship of bacterial bioluminescence contained Chlorophyll a followed by temperature factors with AIC_c_ values of 101.16 (AIC_weight_: 0.50) and 101.95 (AIC_weight_: 0.34). The positive correlation of luminous bacteria with sea surface temperature is in agreement with other results (Fukui et al. 2010, Johnson et al. 2010, Baker-Austin et al. 2016, Montánchez et al. 2019). While inverse relationships of plankton represented by Chlorophyll a with bacterial bioluminescence were also have been reported. Some plankton species were known can release anti bacterial substances that can limit bacteria growth. Taufik et al. (1996) has confirmed that *Thalassiosira* plankton can reduce the density of luminous *V. harveyi* from 8.9 × 10^4^ to 4.3 × 10^4^. This inverse effect of *Thalassiosira* on *V. harveyi* explains low bacterial bioluminescence when Chlorophyll a was high.

## Conclusion

Milky sea and bacterial bioluminescence in Java Southern Sea were presented for 9 days. The size of milky sea and bacterial bioluminescence presence were limited by the low SST and high Chlorophyll a.

